# H5N1 Influenza A is now promiscuous in host range and has improved replication in mammals

**DOI:** 10.1101/2025.03.15.641219

**Authors:** Sayal Guirales-Medrano, Kary Ocaña, Khaled Obeid, Rachel Alexander, Colby T. Ford, Daniel Janies

## Abstract

Influenza A virus has been circulating in birds from Eurasia for more than 146 years, but human infection has been sporadic. H5N1 (clade 2.3.4.4b) has recently infected hundreds of species of wild and domestic birds and mammals in North America. Infections include 70 people with two fatalities. We have developed an analytical bioinformatics, genomics, and structural workflow to understand better how H5N1 is circulating in North America and adapting to new host species.Our time-series analysis reveals that the circulation of H5N1 (clade 2.3.4.4b) in North America follows a distinct annual pattern, with cases in the United States consistently peaking each December. Separate from this seasonal cycle, our analysis also documents an increase in the total number of cases reported since 2021. We also show that H5N1 (clade 2.3.4.4b) spreads in North America as two distinct subclades of interest for human and animal health. These viral lineages have achieved a vast host range by efficiently binding the viral surface protein Hemagglutinin to both mammalian and avian cell surface receptors. This novel promiscuity of host range is concomitant with the additional strengthening of the Polymerase basic 2 viral proteins’ binding for mammalian and avian immune proteins. Once bound, the immune proteins will have diminished ability to fight the virus, thus allowing for more efficient replication of H5N1 in mammalian and avian cells than seen in the recent past. Finally, structural docking analyses predict that while most current antivirals remain effective, a fatal human isolate showed significantly reduced binding to multiple drugs from different classes. In conclusion, the H5N1 virus is causing an animal pandemic through promiscuity of host rage and strengthening ability to evade the innate immune systems of both mammalian and avian cells.

## Introduction

Influenza A viruses (IAVs) are members of the Orthomyxoviridae family, characterized by a genome of eight single-stranded, negative-sense RNA segments. This segmented genome is a critical feature, as it facilitates genetic reassortment, a process where viruses co-infecting a single host can swap RNA segments, leading to the rapid emergence of novel viral strains (1). IAVs are classified into subtypes based on the antigenicity of their two primary surface glycoproteins: hemagglutinin (HA) and neuraminidase (NA). There are 18 known HA subtypes and 11 NA subtypes, which can combine in numerous ways (e.g., H1N1, H3N2, H5N1) (2). Wild aquatic birds, particularly those in the orders of Anseriformes (ducks, geese) and Charadriiformes (gulls, shorebirds), serve as the vast natural reservoir for nearly all IAV subtypes (3). While these viruses typically cause asymptomatic or mild enteric infections in their natural hosts, their introduction into other species can have devastating consequences.

The history of human civilization is marked by recurrent influenza pandemics, which have been exclusively caused by IAVs originating from avian or swine reservoirs. The most catastrophic of these was the 1918 H1N1 “Spanish Flu,” which resulted in an estimated 50 million deaths worldwide (4). Subsequent pandemics, including the 1957 H2N2 “Asian Flu,” the 1968 H3N2 “Hong Kong Flu,” and the 2009 H1N1 swine-origin pandemic, have similarly emerged from the animal-human interface, underscoring the perpetual threat posed by zoonotic influenza (5, 6). This pandemic potential is driven by the continual evolution of the virus through two primary mechanisms: antigenic drift, the gradual accumulation of point mutations, and antigenic shift, the abrupt change caused by genetic reassortment (1).

Among the various avian influenza subtypes, H5N1 has garnered significant global concern. A highly pathogenic avian influenza (HPAI) H5N1 virus first emerged in 1996 in farmed geese in Guangdong, China, belonging to the Goose/Guangdong (Gs/GD) lineage (7). This lineage demonstrated its potent zoonotic capability in 1997 when it caused a severe respiratory illness in 18 people in Hong Kong, resulting in six deaths (8). Since then, HPAI H5N1 has spread across Asia, Europe, Africa, and, more recently, the Americas, evolving into a complex series of genetic clades and causing massive outbreaks in poultry and wild birds (9). The virus has also been responsible for over 985 human infections with a case-fatality at almost 50%, although sustained human-to-human transmission has not been established (10). The ongoing evolution and geographic expansion of HPAI H5N1, particularly its increasing propensity for spillover into mammalian species, necessitates continuous and detailed surveillance.

Alongside vaccination, antiviral medications are a cornerstone of clinical management and pandemic preparedness for influenza. Two primary classes of drugs are licensed in most countries: neuraminidase inhibitors (NAIs), such as oseltamivir and zanamivir, which prevent viral egress from infected cells, and polymerase inhibitors, like baloxavir and marboxil, which targets the polymerase acidic protein (PA) subunit to block viral gene transcription (11). However, the high mutation rate of IAVs presents a constant threat to the efficacy of these drugs. Amino acid substitutions within the viral target proteins can confer resistance, diminishing therapeutic options. For H5N1, which is known to evolve rapidly, it is crucial to monitor mutations in NA, PA, and other polymerase components to ensure that these vital medical countermeasures remain effective (12).

Avian influenza lineages have circulated in Eurasia since at least 1878 (13). We provide a complete account of the history of H5N1’s geographic spread in our recent paper (14). In December 2021, H5N1 Influenza A virus (clade 2.3.4.4b) infection was detected in wild birds in the United States (15). Since then, this viral lineage has spread across the United States and has been detected in a majority of US states (supplemental_fig_30.png). This viral lineage has infected over 246 species (16), including many mammals (17), wild birds (18), cows, cats, and poultry in agricultural settings, and humans coming into contact with wild and farm animals (supplemental_table_1.csv).

As of February 26, 2025, there are 70 confirmed human cases of H5N1 in the United States (19). One death was reported of a 65-year old farm worker in Louisiana, USA, by an H5N1 viral infection of the genotype D1.1 (20). Furthermore, tens of millions of egg-laying chickens have died or been culled in the United States due to H5N1, leading to soaring egg prices (21). In Canada, H5N1 has spread to 132 species of birds and mammals [supplemental_table_2.csv; (22, 23)]. For Mexico, the few species reported by The Servicio Nacional de Sanidad Inocuidad y Calidad Agroalimentaria (SENASICA) are mostly agricultural Galliformes and Anseriformes, with some wild Anseriformes and Pelecaniformes (24). All avian species in SENASICA reports are redundant with species in reports from the USA and Canada (22, 25). The Pan American Health Organization reports no mammal cases in Mexico as of March 4, 2025. However, on April 8, 2025, a young child in Durango, Mexico died due to H5N1 infection of genotype D1.1 (16, 26).

We are seeing a clear increase in the number of reported cases of H5N1 in mammals and avian hosts in the United States (Figure 1). Levels of viral infection have followed seasonal trends where both social and behavioral changes in humans and animals have led to an increase in infection cases (27). H5N1 clade 2.3.4.4b, while present throughout the year, follows the trend of increased infection cases in the winter (primarily December) for temperate climate regions (27). Since the spring and summer of 2022, an increase in summer cases, in North America, has led to a change in the typical trend of H5N1 (28).Through our analysis, we have shown that the seasonality trend associated with the cases of H5N1 in both avian and mammalian hosts in the United States continues to stick with expect increase in winter months (Figure 2).

**Fig. 1.**
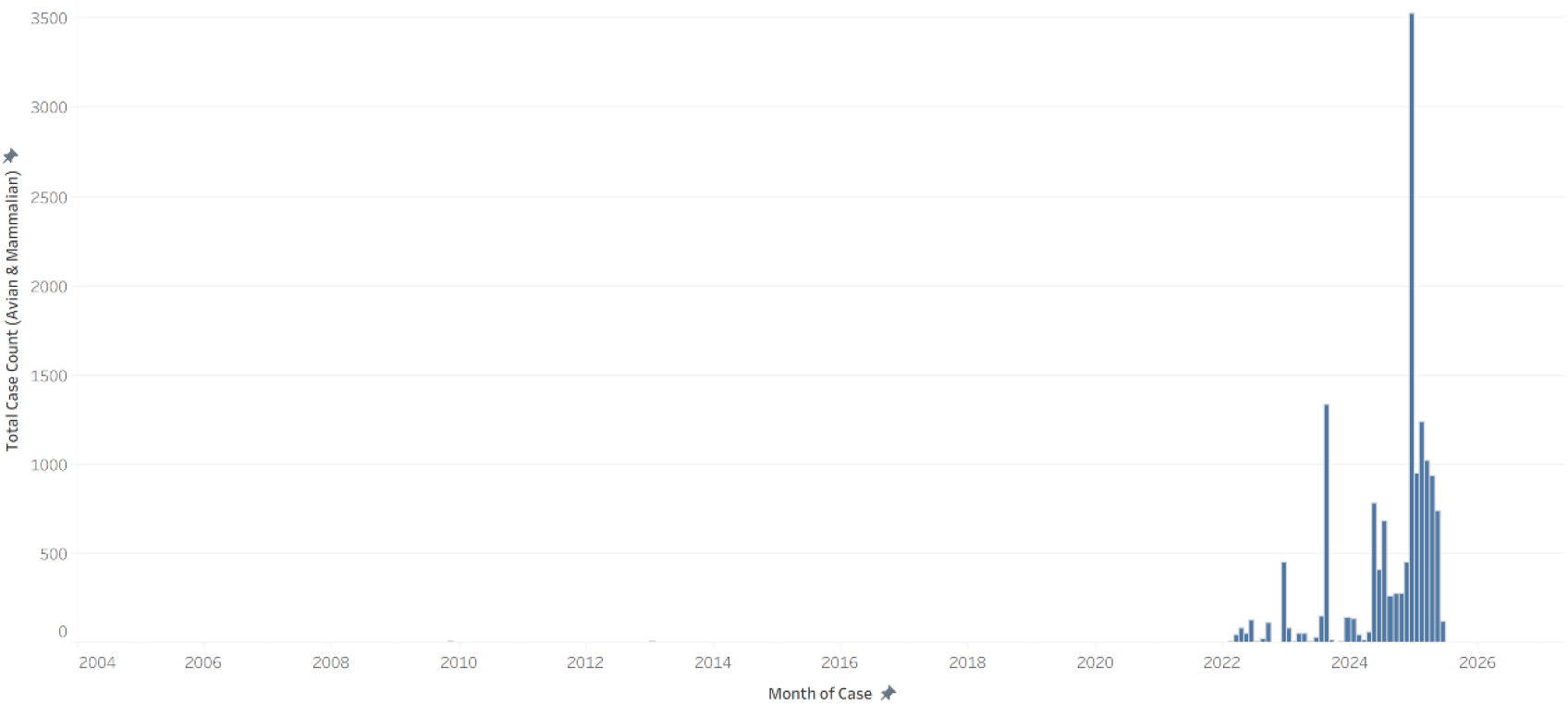
Plot of distinct count of recorded isolates of Influenza A subtype H5N1 versus submission date of isolate as recorded and submitted to GISAID for the United States. These isolates include all isolates reported for mammalian hosts, which consists of “canine, dairy cow, equine, feline, other mammals, [and] swine,” as well as avian hosts, which consists of “chicken, curlew, duck, eagle, falcon, goose, grouse, guineafowl, gull, ostrich, other avian, partridge, passerine, penguin, pheasant, pigeon, rails, sandpiper, shearwater, swan, turkey, turnstone, [and] quail,” as per the GISAID EpiFlue Database description. In total, there were 14,688 unique isolates recorded and submitted, with the highest peak appearing in December 2024 with 3521 recorded isolates. In 2025, there were 1,241 recorded isolates in February, and 114 recorded in June.

**Fig. 2.**
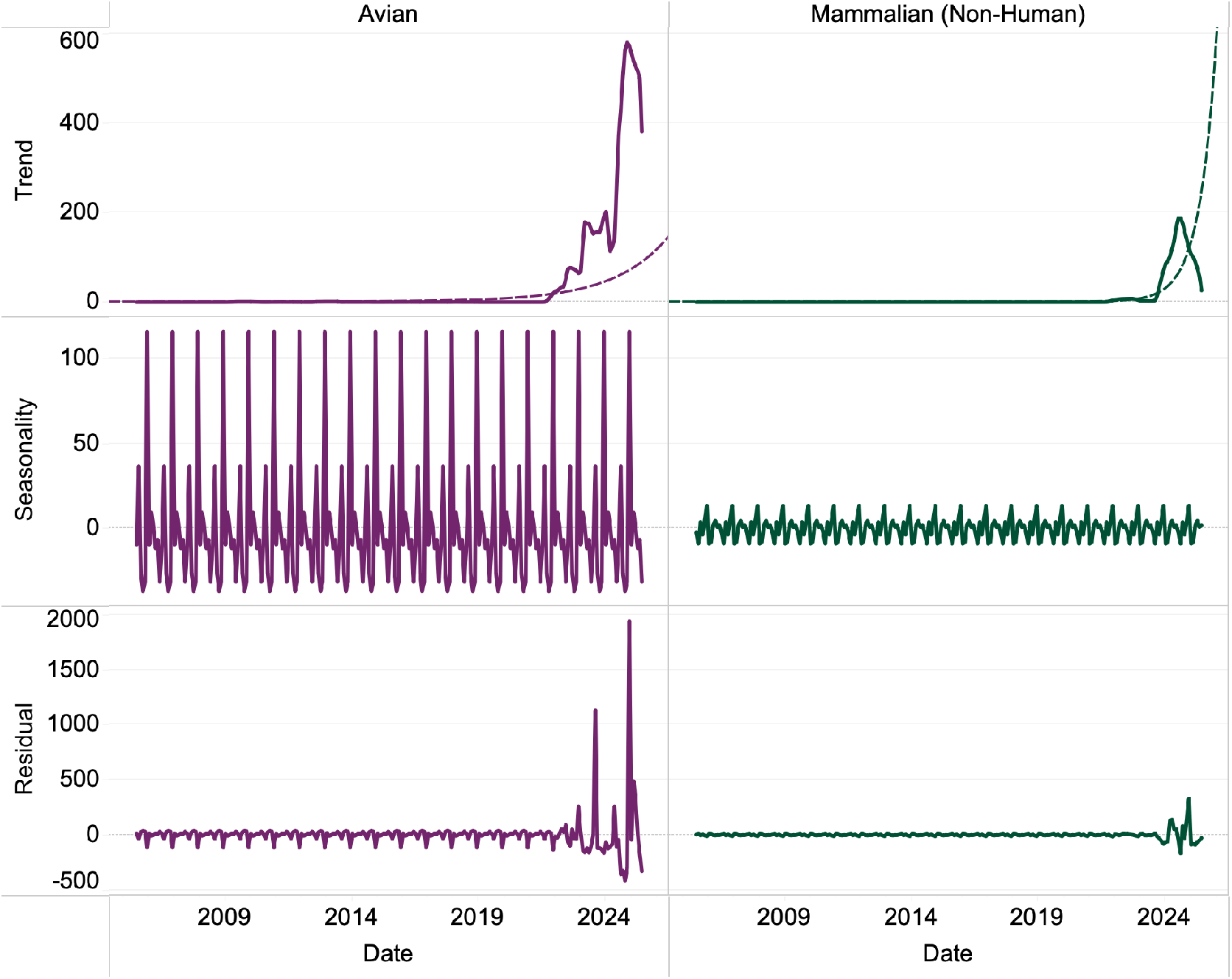
The graph shows seasonal decomposition on United States H5N1 isolates. Data is divided into two columns: avian and mammalian (non-human) isolates. Rows, from top to bottom, show decompositional trend, the residuals, and final seasonality on the y-axis. The x-axis shows time spanning 2008 to 2025. Trend and Residual spikes can be seen occurring starting in 2021. Seasonality shows consistent spikes occurring in December for the United States.

**Fig. 3.**
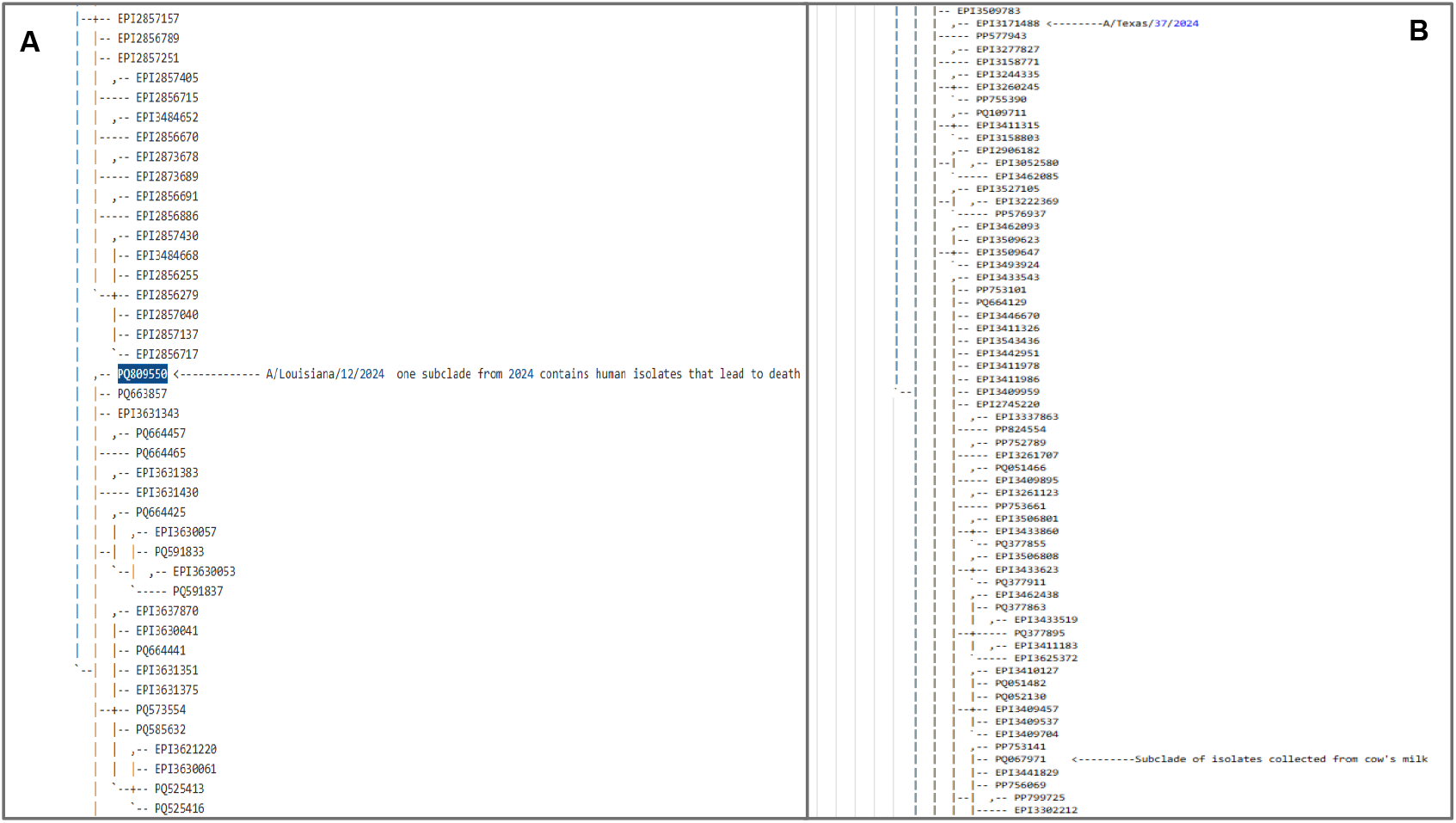
A phylogenetic tree was built from nucleotide sequences of HA using a TNT. ASCII representation of tree is visualized (supplemental_data_file_10.log) within the figure and subclades containing sequences of interest were examined more closely. A) Subclade containing the isolate for the fatal human case in Louisiana. B) Subclade of several cow to human transmission isolates.

Here, we have developed a workflow to study the decomposition of seasonal trends in infection cases, the phylogenetics of the host shifts, natural selection, and structural biology of recent lineages of the H5N1 subtype of influenza A virus in North America. We used the workflow to isolate clades of interest for human infection and study those clades for adaptation to mammals. We assess adaptation using computational analyses of natural selection of the nucleotide diversity of the HA, NA, MP, PA, PB1 and PB2 genes. Once codons under natural selection are identified, we evaluate the nature of the adaptation of H5N1 (clade 2.3.4.4b) viral lineages through computational structural biology. Here, we show that Hemagglutinin (HA) mutations have shifted the virus from adaptation to avian cells to promiscuity to many species of wild and domestic mammals and birds. Our analyses of natural selection in Polymerase Basic 2 (PB2) show that mutations have adapted the H5N1 (clade 2.3.4.4b) viral lineage to increase their ability to bind proteins important to the viral invasion and immune response of the avian and mammalian host cells, thus dampening the hosts’ innate defenses to infection and setting the stage for increased viral replication. Finally, our analyses on the structural changes of recent antiviral target proteins Neuraminidase (NA), Matrix protein 2 (MP), Polymerase Acid (PA), PB1 and PB2, have shown continued effect of some antiviral treatments but potentially worsened efficacy of others.

## Methods

### Assessment of Seasonality

#### Case Collection

Data to assess seasonality were collected from the Global Initiative on Sharing All Influenza Data (GISAID; gisaid.org) EpiFlu Nucleotide Sequence Database. Filters for subtypes H5 and N1 were applied, in addition to host and location. Hosts were analyzed as avian and mammalian (non-human). Species included as avian hosts, per GISAID, are as follows: “chicken, curlew, duck, eagle, falcon, goose, grouse, guineafowl, gull, ostrich, other avian, partridge, passerine, penguin, pheasant, pigeon, rails, sandpiper, shearwater, swan, turkey, turnstone, [and] US quail.” Species included as mammalian hosts, per GISAID, are as follows: “canine, dairy cow, equine, feline, other mammals, [and] swine.” Localities were filtered to be either North America, outside North America, or exclusively within the United States. The data was then tabulated based on the collection date of each viral isolate.

#### Data Visualization

Isolate data was visualized through creating a series of dashboards on Tableau (https://www.tableau.com), in which viral isolates grouped by host and location were graphed versus time. Viral isolate data was collected from several sources, including GISAID’s Nucleotide Sequence Database, the United States Department of Agriculture’s Animal and Plant Health Inspection Service (USDA, APHIS), the World Animal Health Information System’s (WAHIS) Disease Data Collection, and the World Health Organization (WHO). After aggregation, these data were processed into groups for host and for location. Groups for hosts included: avian (wild and agricultural), mammalian (wild and agricultural, non-human), and human.

#### Seasonality Decomposition

The time series case data was further analyzed through seasonal decomposition. Using the seasonal_decompose() method from the statsmodels Python library, a decomposition of the time series using moving averages resulted in separate time series that described the trend, the seasonality, and the residual (error) of the original data (29).

### Data Acquisition and Preparation

#### Sequence Dataset Compilation

The H5N1 Hemagglutinin (HA) dataset from Ford et al. (2025) (14) was updated with sequences released in the latter half of 2024 and early 2025, sourced from GISAID’s EpiFlu database and the United States National Institutes of Health’s Genbank (NCBI; ncbi.nih.nlm.gov) database. The initial dataset comprised 18,289 HA sequences. A second dataset of Polymerase Basic 2 (PB2) nucleotide sequences from clade 2.3.4.4b and relevant background strains was similarly compiled from GISAID and GenBank (30–32). Additionally, a dataset of 29,011 nucleotide sequences were collected for MP, 18,466 nucleotide sequences for NA, 17,873 nucleotide sequences for PA, and 14,089 nucleotide sequences for PB1.

#### Multiple Sequence Alignment and Curation

From the comprehensive HA dataset, a working set of 9,359 sequences representing clade 2.3.4.4b and background data was selected (supplemental_table_3.csv). These sequences were realigned using MAFFT (33). For each gene dataset, alignments were manually curated by trimming terminal unaligned regions (“ragged edges”), padding any remaining terminal gaps with ‘?’, and removing any sequences that disrupted the translational reading frame. After this curation, the final HA dataset consisted of 9,359 sequences (supplemental_data_file_4.nexus). We spliced the nucleotides from M1 and M2 to the reading frame to make a coding region for MP. The final MP dataset consisted of 22,858 sequences (supplemental_data_file_18.nexus). The final NA dataset consisted of 14,000 sequences which were then reduced further into 3 subclades. Downstream analyses used subclade 3 consisting of 257 sequences (supplemental_data_file_19.nexus). The final PA dataset consisted of 429 sequences (supplemental_data_file_20.nexus). The final PB1 dataset contained 9,841 sequences (supplemental_data_file_21.nexus). The final PB2 dataset contained 3,415 sequences aligned to 2,281 positions (supplemental_data_file_5.nexus). Alignments were numbered sequentially following realignment.

### Phylogenetic and Evolutionary Analysis

#### Large-Scale Phylogenetic Tree Construction

Initial phylogenetic trees for the full HA, NA, MP, PA, PB1 and PB2 datasets were constructed using TNT (Version 1.6; (34)) with equal weighting for character state changes. GISAID sequence EPI100512 (A/chicken/Kulon Progo/BBVet-XII-1/2004) was used as the outgroup for the HA analysis, and NCBI sequences GU052520, GU052519, GU052523, GU052524, GU052525 was used for the NA, MP, PB1 and PB2 analyses respectively (A/chicken/Scotland/1959).

#### Character Evolution Mapping

For the full HA dataset (9,359 sequences), two independent characters (host and date of isolation) were defined based on sequence metadata (supplemental_table_3.csv). The evolutionary history of these characters was mapped onto the HA phylogeny using Mesquite to trace host and temporal transitions within clades of interest (35). Similar character mapping was performed on the other gene phylogenies centering around clades formed by the same isolates of interest.

### Phylogenetic and Network Analysis

For HA, NA, and PB2 we planned downstream structural analyses and thus focused on recent subclades containing human isolates and subclades containing recent zoonotic strains.

- HA Subclade 1 (surrounding fatal Louisiana case): This dataset focused on isolates phylogenetically related to a human case in Louisiana, comprising 331 sequences aligned to 1,701 positions (supplemental_data_file_6.phy).
- HA Subclade 2 (human-bovine-avian events): This dataset focused on transmission events between human, bovine, and avian hosts, comprising 767 sequences aligned to 1,701 positions (supplemental_data_file_7.phy).
- PB2 Subclade (2022-2025 zoonotic events): A subclade subtended by strain CY205506 was selected, containing 1,528 sequences aligned to 2,277 positions (supplemental_data_file_8.phy).
- NA Subclade 3 (2022-2025 zoonotic events): A subclade containing 257 sequences aligned to 1,410 positions.

For MP, PA, and PB1, which were not subject to downstream structural analyses, we used full longitudinal datasets.

For all datasets, stop codons were trimmed to prepare for selection analyses. Phylogenetic trees for all subclades were inferred using RAxML (v8.2.12; (36)) under GTRGAMMA. For HA, Genbank sequence LC718258 (A/jungle crow/Hokkaido/0104B085/2022) and GISAID sequence EPI3509783 (A/chicken/Montana/24-001136-002-original/2024) served as outgroups for Subclade 1 and 2, respectively. For NA we used EPI460427 (A/turkey/Ontario/6213/1966) as an outgroup. For MP we used NC_00736 (Goose/Guangdong/1996) as an outgroup. For PA we used GU052523 (A/chicken/Scotland/1959) as an outgroup. For PB1 we used GU052524 (A/chicken/Scotland/1959) as an outgroup.

StrainHub networks were then generated for each HA subclade to visualize transmission dynamics using betweenness centrality (Figures 4 and 5) (37).

**Fig. 4.**
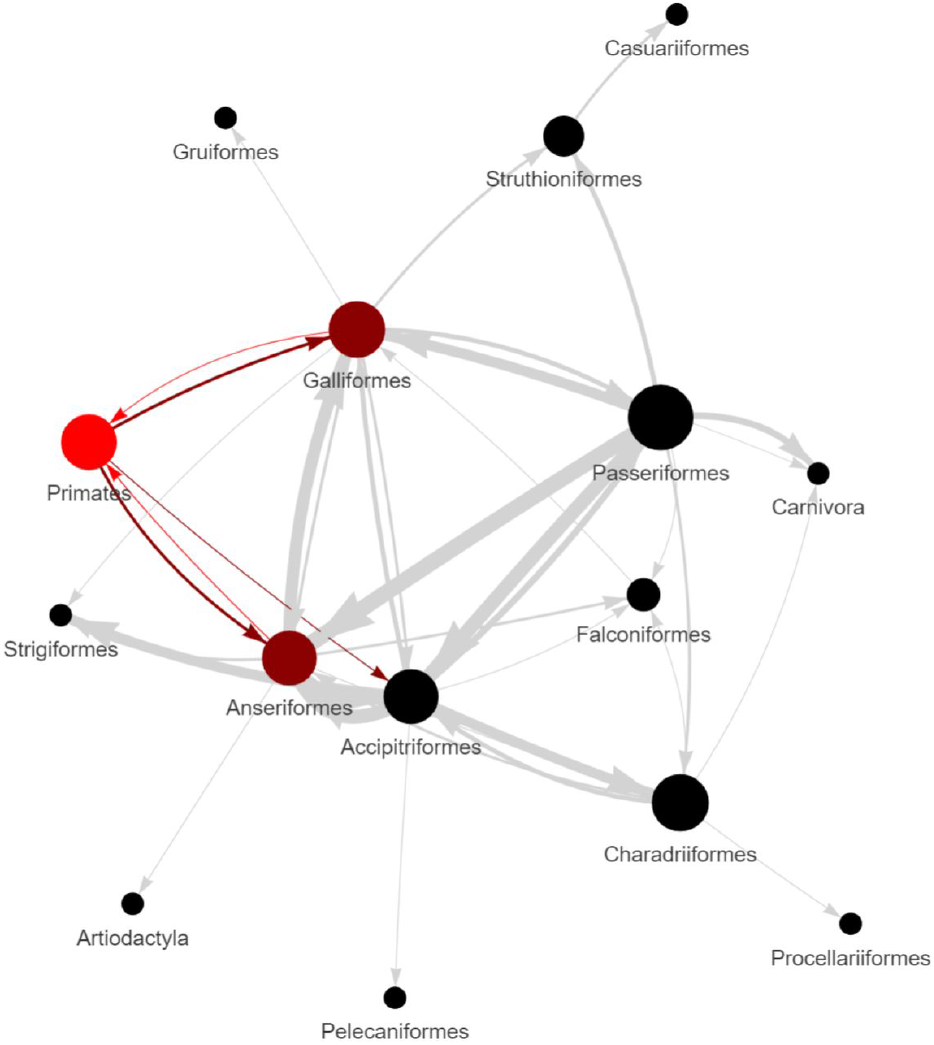
A Strainhub network using a tree based on HA sequence data and host metadata for the clade of interest surrounding the deceased patient infected with H5N1 in Louisiana in 2024 (PQ809550). The metric used to create the network was betweenness centrality. The arrows represent the directionality of viral transmission events recovered. The thickness of the lines represents a higher frequency of transmission. The colors represent highlighted zoonotic events such as bidirectional transmission between Primates (humans), Galliformes (e.g., poultry), and Anseriformes (e.g., ducks and geese), and unidirectional transmission from Primates to Accipitriformes (e.g., hawks and eagles).

**Fig. 5.**
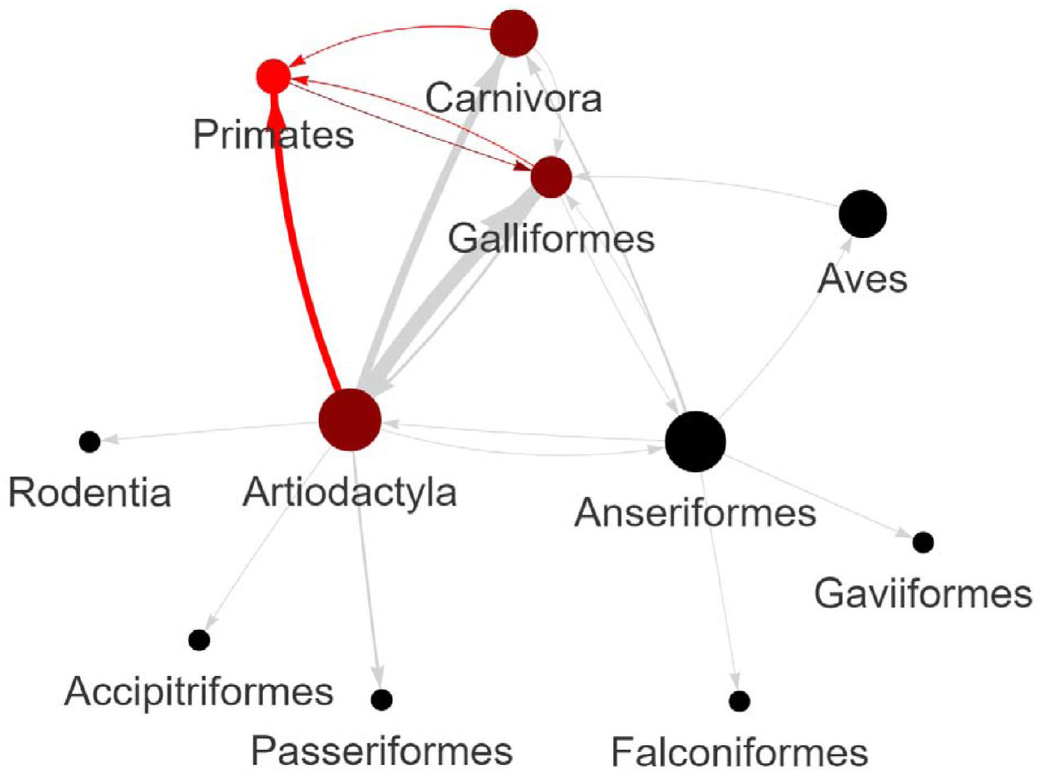
A Strainhub network using a tree based on HA sequence data and host metadata for the clade of interest surrounding the human-bovine-avian transmission events in 2024. The metric used to create the network was betweenness centrality. The arrows represent the directionality of viral transmission events recovered. The thickness of the lines represents a higher frequency of transmission. The colors represent highlighted zoonotic events such as unidirectional transmission from Artiodactyla (e.g., cows) to Primates (humans), bidirectional transmission between Primates and Galliformes (e.g., poultry), and unidirectional transmission from Carnivores (e.g. domestic cats) and Primates.

### Natural Selection Analysis

Prior to analysis, all subclade datasets were inspected for internal stop codons using MACSE (v2) and manual review (38). Candidate vaccine sequences (e.g., GISAID: EPI1846961 for HA; Genbank: OQ958046 for NA, OQ958047 for MP, OQ958043 for PA, OQ958058 for PB1, OQ958041 for PB2) were added as reference baselines. In preparation for HYPHY we calculated trees using RAxML under GTRGAMMA. Analyses of natural selection were performed using the HyPhy package (v2.5.67; (39)). We employed the Nielsen and Yang (1998) approach to estimate the ratio of nonsynonymous to synonymous substitution rates (dN/dS) per codon (40). The posterior probabilities of a site being under positive selection (dN/dS>1) were calculated. A suite of models was tested to ensure a robust evaluation of evolutionary pressures: M0 (Single Rate), M1 (Neutral), M2 (Selection), M3 (Discrete), M7 (Beta), M8 (Beta & ω), and M12 (Two-Class). This allowed for the detection of both pervasive purifying selection and episodic positive selection across the datasets.

### Structural Biology and Molecular Docking

#### Hemagglutinin (HA) Structural Analysis

Three-dimensional structures were generated for four HA protein sequences of interest: GISAID: EPI3171488, EPI1846961 (vaccine candidate), Genbank: PQ591824, and PQ809550. Amino acid sequences were folded using ColabFold (v1.5.2; (41)). To simulate interactions with host receptors, human and avian Sialic Acid (SA) glycan structures were obtained from PDB entries 4K63 and 4KDO, respectively. Protein-glycan docking was performed for each HA model against both human and avian SA glycans using HADDOCK3 (v2024.12.0b7; (42)) to assess potential differences in binding affinity.

#### Polymerase Basic 2 (PB2) Structural Analysis

A similar structural analysis was conducted for the PB2 protein. Sequences from strains OQ958041 (vaccine candidate), PQ809562, PQ591825, and PP577947 were selected. Additionally, sequences carrying mutations at position 588 [GISAID: EPI3304108 (A588T), EPI3315580 (A588S), and EPI3447228 (A588V)] were included. Known host interaction partners of PB2 were selected as docking targets: human MAVS (PDB: 2MS8), human Importin-*α*3 (PDB: 4UAE), its avian ortholog (UniProt: A0A2I0M1B5), human ANP32A (PDB: 6XZQ), and its avian ortholog (NCBI: XP_075287452). Each protein target selected showcases a different interaction with influenza’s PB2. MAVS is responsible for innate immune response signaling of JAK1 which is inhibited by PB2 interactions (43). Importin-*α*3 promotes transport of proteins into the host nucleus, i.e. importing PB2 into the nucleus for viral replication (44). Lastly, ANP32A was selected due to its ability to mediate and increase viral replication when interacting with PB2 (45). All protein structures were folded using ColabFold. Protein-protein docking simulations between PB2 variants and host targets were conducted using HADDOCK3 to evaluate binding interactions.

#### Antiviral Docking Analysis

To investigate potential variations in antiviral susceptibility, a structural docking analysis was performed on key H5N1 protein targets. The selected proteins included neuraminidase (NA), a surface glycoprotein that mediates viral egress, and the three components of the RNA-dependent RNA polymerase complex: PA, PB1, and PB2. Sequences for these proteins were obtained from NCBI and GISAID for a panel of representative strains:

- A/chicken/Scotland/1959 (historical reference)
- A/American Wigeon/South Carolina/22-000345-001/2021 (vaccine candidate)
- A/Astrakhan/3212/2020 (vaccine candidate)
- A/cattle/Texas/24-009110-004/2024 (bovine-to-human transmission)
- A/Texas/37/2024 (bovine-to-human transmission)
- A/California/150/2024 (bovine-to-human transmission)
- A/Louisiana/12/2024 (fatal avian-to-human transmission)

The corresponding antiviral compounds were selected based on their known protein targets: Oseltamivir and Zanamivir (NA inhibitors), Baloxavir (PA inhibitor), Favipiravir (PB1 inhibitor), and Pimodivir (PB2 inhibitor). The SMILES (Simplified Molecular-Input Line-Entry System) strings for each antiviral were retrieved from the PubChem database (46).

The computational workflow for this analysis was as follows: The 3D structure of each protein variant was predicted using ColabFold. Concurrently, the SMILES strings for the antiviral compounds were converted into 3D molecular structures using RDKit (47). Putative binding pockets on each target protein were identified using P2Rank (48). Finally, protein-ligand docking simulations were conducted between each antiviral and its respective target protein using HADDOCK3 to evaluate their binding interactions.

## Results

### Rise of H5N1 and the question of seasonality of outbreaks in the United States

In order to assess seasonality of outbreaks we focused on non-human cases recorded in GISAID. Through a graph of isolates of H5N1 from the United States in GISAID over time, we see a large increase in mammalian (non-human) and avian cases in late 2024 as shown in Figure 1.

In response to the acceleration (peaking in winter 2024) and deceleration (waning in the spring and summer of 2025) of H5N1’s outbreaks in the United States questions have arisen as to whether the virus has been controlled or is it in a seasonal lull (49). To investigate the temporal dynamics of H5N1 outbreaks, we performed a time-series analysis on reported isolates from avian and mammalian (non-human) hosts in the United States (Figure 2). The analysis revealed a dramatic, statistically-significant exponential increase in the total number of reported H5N1 isolates, with this surge beginning in 2021. The trend plots clearly indicate that this recent spike in recorded isolates is predominantly driven by outbreaks in avian hosts. Further analysis of the seasonality and frequency with which viral isolates were collected indicates that there was a significant uptick of outbreak activity in both avian and mammalian (non-human) hosts in December 2024.

In fact, reporting on GISAID showed that there were 449 recorded isolates in November 2024 and 3,521 recorded isolates in December 2024, indicating a nearly 684% increase in confirmed cases for avian and mammalian (non-human) hosts in the United States. This trend is similarly reported by the USDA APHIS, where there were 552 confirmed cases in November 2024 and 1,189 confirmed cases in December 2024, showing an increase by 115% (50).

Our analysis identified a seasonal pattern of outbreak intensity within the United States between 2021 and 2025. Cases in both avian and mammalian (non-human) hosts peaked late in the calendar year, typically in December.

### Phylogenetic Analysis Reveals Host-Shifting in North American H5N1

To understand the patterns of host transmission, we analyzed 9,359 unique H5N1 HA sequences from North America in three heuristically parsimonious trees were found at 16950 steps using TNT (supplemental_data_file_9.log) (34). We used two of the three independent tree searches. We also mapped host category (Avian, Mammalian) and date of isolation onto the phylogeny. This analysis revealed multiple independent spillover events from the primary avian reservoir into various mammalian species. The temporal mapping showed consistent progression of the virus across the continent over the three-year period.

Based on these patterns, we identified two key subclades for in-depth analysis due to their association with human infections:

1. A clade surrounding a fatal human case in Louisiana ((51);Genbank accession PQ809550) is seen in Figure 3A (supplemental_data_file_10.log). Sequences in this subclade are referred to as genotype D1.1.
2. A separate, large clade involving transmission between wild birds, poultry, cattle, and a human in Texas is seen in Figure 3B (supplemental_data_file_10.log). This subclade contains HA sequences from influenza isolates referred to as genotype B3.13.

Strainhub analysis of the HA subclades of interest show the transmission networks between hosts. In these figures the arrow-heads represent the direction(s) of the transmission events. The thickness of the lines represent the frequency of the transmission events. Colors are for emphasis.

In Figure 4, the subclade that contains the virus that led to the fatal human case in Louisiana shows significant transmission moving bidirectionally from primates (including humans) towards Galliformes and Anseriformes, and back from these two groups towards Primates.

In Figure 5, the human-bovine-avian transmission clade exhibits bidirectional transmission among Galliformes and Primates. There is transmission from Carnivora towards Primates. There is high frequency transmission from Artiodactyla to Primates.

The curated data for PB2 are contained in an alignment of 3415 sequences and 2281 aligned positions (supplemental_data_file_5.nexus). In three independent TNT searches, we found 465 heuristically parsimonious trees of 12133 steps (supplemental_data_file_11.log). We chose the first tree to select a subclade of interest based on the position of isolates of interest (supplemental_data_file_12.log). We identified the subclade subtended by the outgroup CY205506 and removed sequences with stop codons to ready the data for analyses of natural selection. The resulting data contained 1528 taxa and 2277 aligned positions (supplemental_data_file_8.phy).

Curated data from these subclades were carried forward for detailed analyses of natural selection and protein structure.

### Positive Selection on HA and PB2

Table 1 lists the results for the HA subclade of interest for avian-human exchanges, including the lineage that led to the death of the human patient in Louisiana (Figure 4 and supplemental_data_file_6.phy plus EPI1846961, A/Astrakhan/3212/2020 as a baseline). The results for the HA subclade of interest for avian-bovine-human exchanges (Figure 5 and supplemental_data_file_7.phy plus EPI1846961, A/Astrakhan/3212/2020 as a baseline).

**Table 1.** Rows 1 through 5 contain HA amino acid sites (using sequential site numbering) with dN/dS>1 (model 8; posterior cutoff = 0.95) in the clade surrounding the deceased patient infected with H5N1 in Louisiana. In the rightmost column we list the mutations observed in these data. Rows 6 through 10 contain HA amino acid sites (using sequential site numbering) with dN/dS>1 (model 8; posterior cutoff = 0.95) in human-bovine-avian clade. In the rightmost column we list the mutations observed in these data. Rows 11 through 18 contain PB2 amino acid sites (using sequential site numbering) with dN/dS>1 (model 8; Posterior cutoff = 0.95; *The probability of site 588 is very close to the cutoff and thus was considered in structural analyses) in current H5N1 lineages. In the rightmost column we list the mutations observed in these data.

| Protein | Clade | Site | Probability | Mutation(s) |
| --- | --- | --- | --- | --- |
| HA | surrounding the deceased patient infected with H5N1 in Louisiana | 140 | 0.960974714844818 | N140D/H |
| HA | surrounding the deceased patient infected with H5N1 in Louisiana | 230 | 0.9685961144425779 | A230V |
| HA | surrounding the deceased patient infected with H5N1 in Louisiana | 298 | 0.9697143147489277 | V298I/L |
| HA | surrounding the deceased patient infected with H5N1 in Louisiana | 490 | 0.9569366098585621 | D490N |
| HA | surrounding the deceased patient infected with H5N1 in Louisiana | 509 | 0.9682055136291944 | E509G |
| HA | human-bovine-avian | 11 | 0.9733589348171219 | V11I |
| HA | human-bovine-avian | 147 | 0.9988807760818452 | V147M |
| HA | human-bovine-avian | 172 | 0.9602969606372659 | A172T |
| HA | human-bovine-avian | 250 | 0.9680511447623218 | K250E/N/R |
| HA | human-bovine-avian | 526 | 0.9894637570633174 | I526V |
| PB2 | current H5N1 lineages | 184 | 0.9670923982786206 | T184A/M/K |
| PB2 | current H5N1 lineages | 249 | 0.9973568275209534 | E249G/D/K/V |
| PB2 | current H5N1 lineages | 292 | 0.9922498777048648 | I292L/V/T/M |
| PB2 | current H5N1 lineages | 344 | 0.9999928310390789 | V344M/A/I |
| PB2 | current H5N1 lineages | 463 | 0.9789001325108995 | I463M/V |
| PB2 | current H5N1 lineages | 588 | 0.9428168996255263* | A588T/S/V |
| PB2 | current H5N1 lineages | 627 | 0.9999895456621494 | E627K/V |
| PB2 | current H5N1 lineages | 676 | 0.9666441948315898 | T676A/I/L/V |

Table 1 also lists the results for the PB2 subclade of interest (refer to supplemental dataset supplemental_data_file_8.phy plus OQ958041, A/American Wigeon/South Carolina/22-000345-001/2021 as a baseline).

Details on the function of these mutations, if known, are provided in the discussion.

### Positive Selection on Antiviral Target Proteins

Table 2 shows the results of determining positive selection on the subclades identified for the proteins NA (supplemental_data_file_19.nexus plus EPI460427, A/turkey/Ontario/6213/1966 as a baseline), MP (supplemental_data_file_22.phy plus NC_007363, A/goose/Guangdong/1/1996 as a baseline), PA (supplemental_data_file_20.nexus plus GU052523, A/chicken/Scotland/1959 as a baseline), and PB1 (supplemental_data_file_23.phy plus GU052524, A/chicken/Scotland/1959 as a baseline). No amino acid sites were found to be under positive selection for proteins PA and PB1 within our dataset.

**Table 2.**
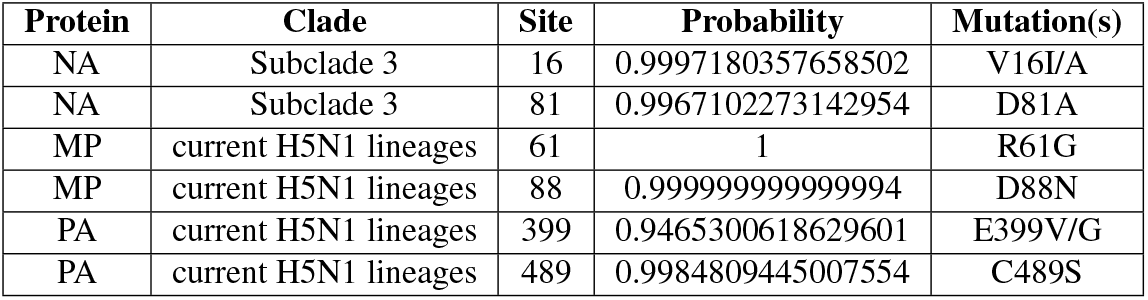
Table shows NA, MP, and PA amino acid sites (using sequential site numbering) with dN/dS>1 (model 8; Posterior cutoff = 0.95). In the rightmost column we list the mutations observed in these data. No amino acid sites were identified to have positive selection in PB1.

| Protein | Clade | Site | Probability | Mutation(s) |
| --- | --- | --- | --- | --- |
| NA | Subclade 3 | 16 | 0.9997180357658502 | V16I/A |
| NA | Subclade 3 | 81 | 0.9967102273142954 | D81A |
| MP | current H5N1 lineages | 61 | 1 | R61G |
| MP | current H5N1 lineages | 88 | 0.9999999999999994 | D88N |
| PA | current H5N1 lineages | 399 | 0.9465300618629601 | E399V/G |
| PA | current H5N1 lineages | 489 | 0.9984809445007554 | C489S |

### Structural Analyses Reveal Increased Affinity for Mammalian Host Factors

To understand the functional consequences of the identified mutations, we performed a series of molecular docking simulations using HADDOCK3 and studied Van der Waals’ energy as the primary measure of binding affinity.

#### Hemagglutinin Docking

Analysis of the HA-SA docking results indicated similar in binding affinity as measured by Van der Waals’ energy between HA and the mammalian (human) or avian SA glycan variants in many clades except the clade containing the fatal human case in Lousiana (PQ809562, A/Louisiana/12/2024 ; Figure 6). In the fatal human case in Louisiana the binding of HA to SA was much stronger in mammalian SA than avian SA (Figure 6 orange bars on the right).

**Fig. 6.**
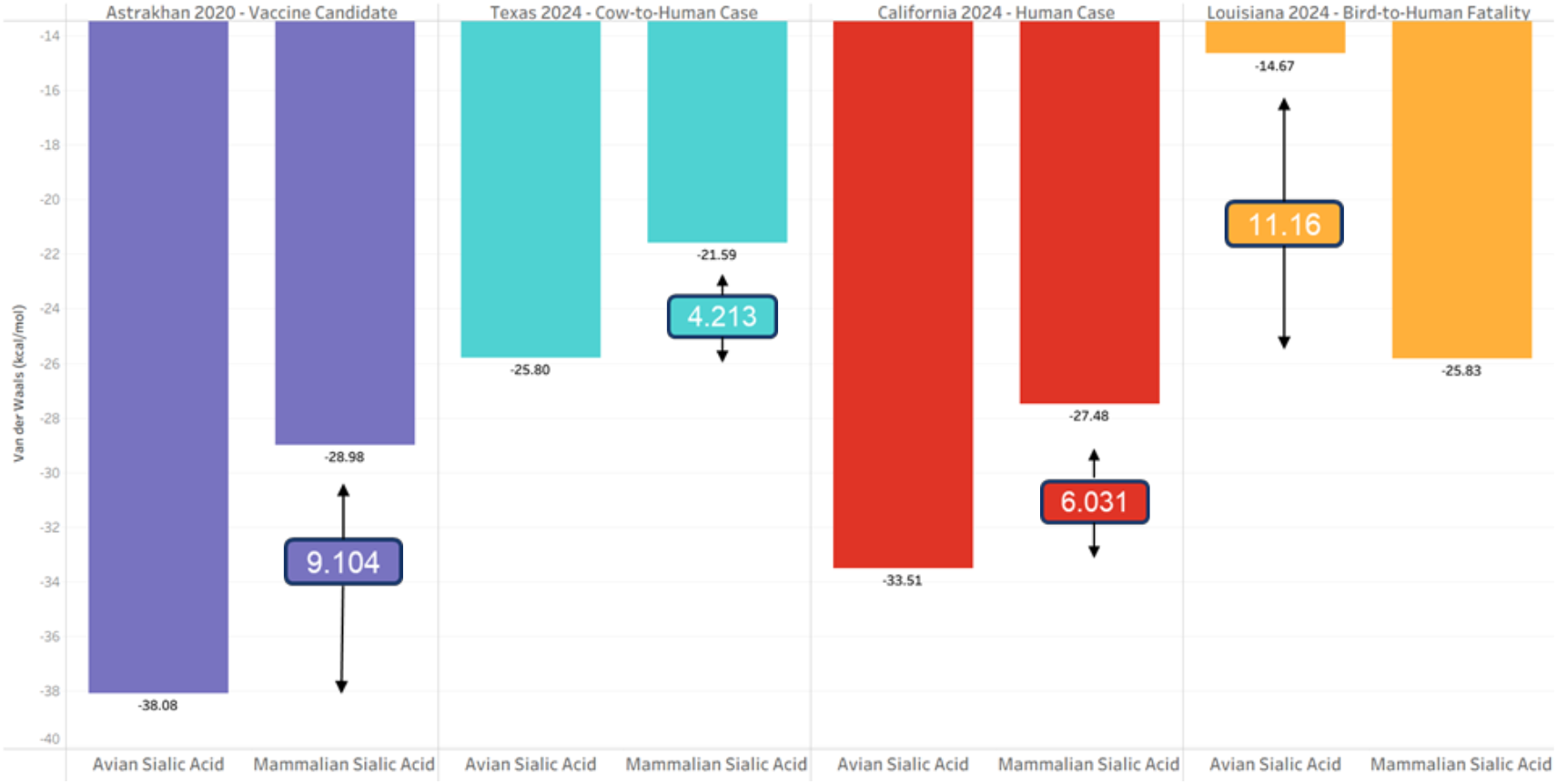
The bar graph shows the results of HADDOCK3 docking simulations for Hemagglutinin (HA) binding to avian and human sialic acid (SA) glycans. The y-axis represents the Van der Waals’ energy (VDW), which measures binding affinity, with lower values indicating stronger binding interactions. The x-axis categorizes the results based on the type of SA (avian or human). Each colored represents an individual sample [blue: EPI1846961 (A/Astrakhan/3212/2020, vaccine candidate), cyan: EPI3171488 (A/Texas/37/2024), red: PQ591824 (A/California/150/2024), and orange: PQ809550 (A/Louisiana/12/2024)]. The numbers boxed between the Avian and Mammalian bars shows the differences between the two VDW energies in kcal/mol. The largest difference is seen in A/Louisiana/12/2024 (orange).

Across the other viral strains (EPI1846961, A/Astrakhan/3212/2020; PP577947, A/Texas/37/2024; PQ591825, A/California/150/2024; Figure 6, blue, cyan, and red bars), HA exhibited comparable binding interactions with both human and avian SA.

Additionally, in supplemental_fig_29.png, we see the spatial orientation and positioning of SA molecules within the docking simulations remained consistent, reinforcing the conclusion that H5N1 clade 2.3.4.4b HA maintains the ability to bind both human and avian SA without structural bias. This suggests that the selected HA variants retain the capacity for a wide host range and adaptation.

#### Polymerase Basic 2 Structures

In contrast to results for HA binding to SA glycans, docking simulations of polymerase PB2 viral protein motifs with host immune response proteins revealed an increased binding affinity in all recent PB2 strains compared to the vaccine candidate (OQ958041, A/American Wigeon/South Carolina/22-000345-001/2021; Figure 7). A/Louisiana/12/2024 is the most immune evasive. These findings indicate an enhanced ability of recent PB2 variants to inhibit host innate immune signaling pathways, contributing to greater immune evasion (52).

**Fig. 7.**
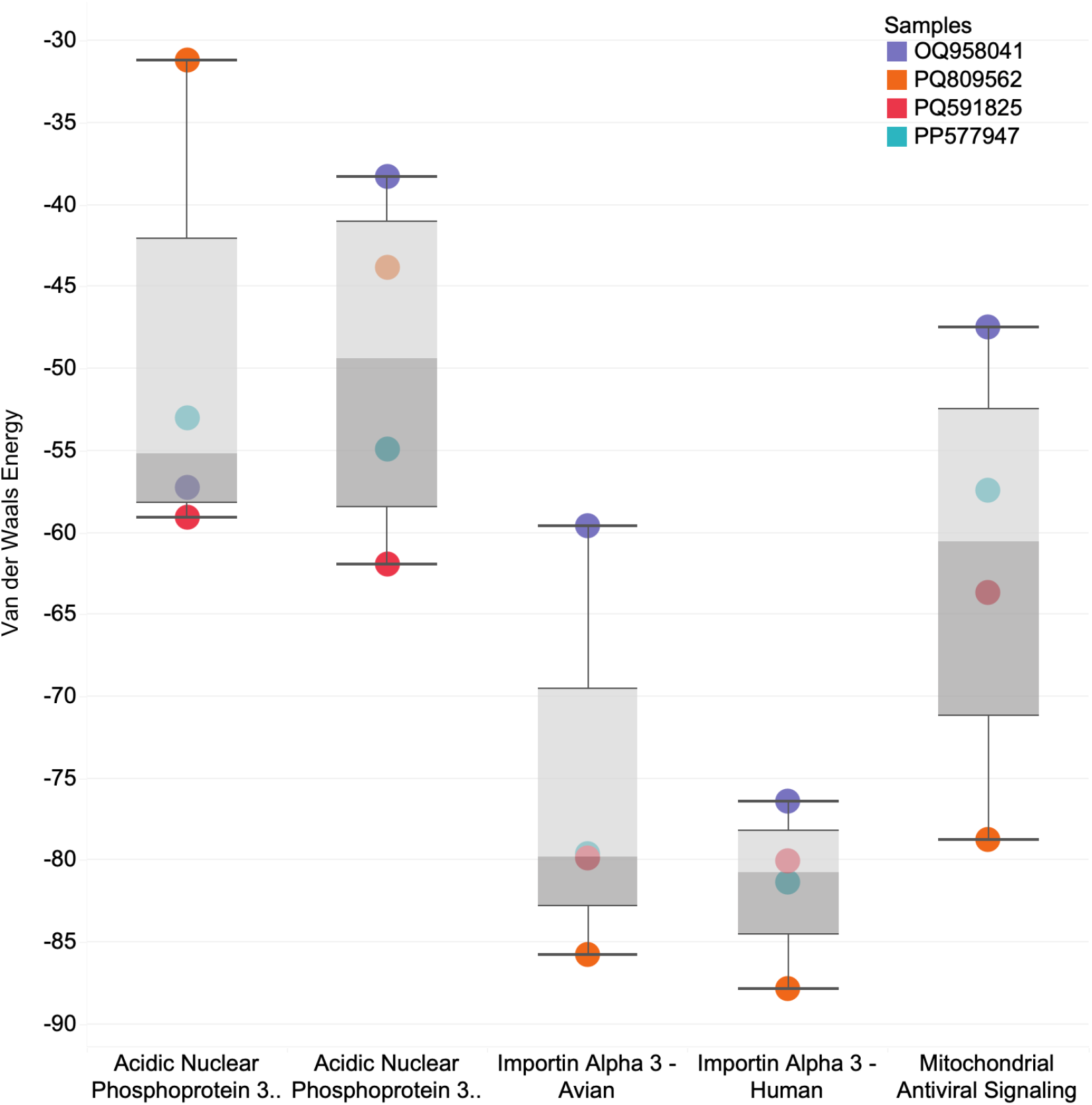
Boxplot graph showing the HADDOCK3 docking results for Polymerase Basic 2 (PB2) binding to various host immune response proteins. The y-axis represents the Van der Waals’ energy (kcal/mol). We use this metric to assess binding affinity, with lower values indicating stronger interactions. The x-axis categorizes the results based on the target proteins: ANP32A (avian and human variants), Importin-*α*3 (KPN4; avian and human variants), and MAVS. Each colored point corresponds to an individual PB2 viral protein [blue: OQ958041 (A/American Wigeon/South Carolina/22-000345-001/2021, vaccine candidate), orange: PQ809562 (A/Louisiana/12/2024),red: PQ591825 (A/California/150/2024), and cyan: PP577947(A/Texas/37/2024)].

The results indicate that PB2 exhibits differential binding affinities to its target host proteins. Notably, some recent PB2 strains show stronger binding interactions (lower Van der Waals’ energy) than the vaccine candidate (OQ958041), for Importin-*α*3 (KPN4) and MAVS. We also see improved binding to human ANP32A as compared to avian ANP32A, which indicates improved replication within human cells (45). This trend indicates strengthening immune evasion capabilities among recent PB2 variants in both human and avian hosts. In summary, binding experiments with viral and host proteins across both North American H5N1 clades of interest show a clear increase in affinity for mitochondrial antiviral signaling proteins in human and mammalian infections, along with enhanced viral replication and nuclear localization in both human and avian hosts.

#### Binding of viral proteins carrying mutations at PB2 protein site 588, including GISAID: EPI3304108 (A588T mutation), EPI3315580 (A588S mutation) EPI3447228 (A588V mutation)

Improved binding for ANP32A and MAVS remains the same in A588T/S/V variants (i.e. EPI3304108, EPI3315580, and EPI3447228) as isolates representing clades of interest mentioned above (OQ958041, PQ809562, PQ591825, and PP577947). The 588 mutation occurs in the ‘627 domain’ which we test by binding to KPN4 (53). Most isolates with variation at site 588 (A588 have an improved binding affinity in humans than in avian host proteins except for the A588S mutation as seen in Figure 9. A588S caused a decrease in affinity to human proteins. For A588S, we also see that binding affinity is worse than the vaccine candidate in the human KPN4. No reported effects of the 588S mutation in PB2 have been reported in literature.

**Fig. 8.**
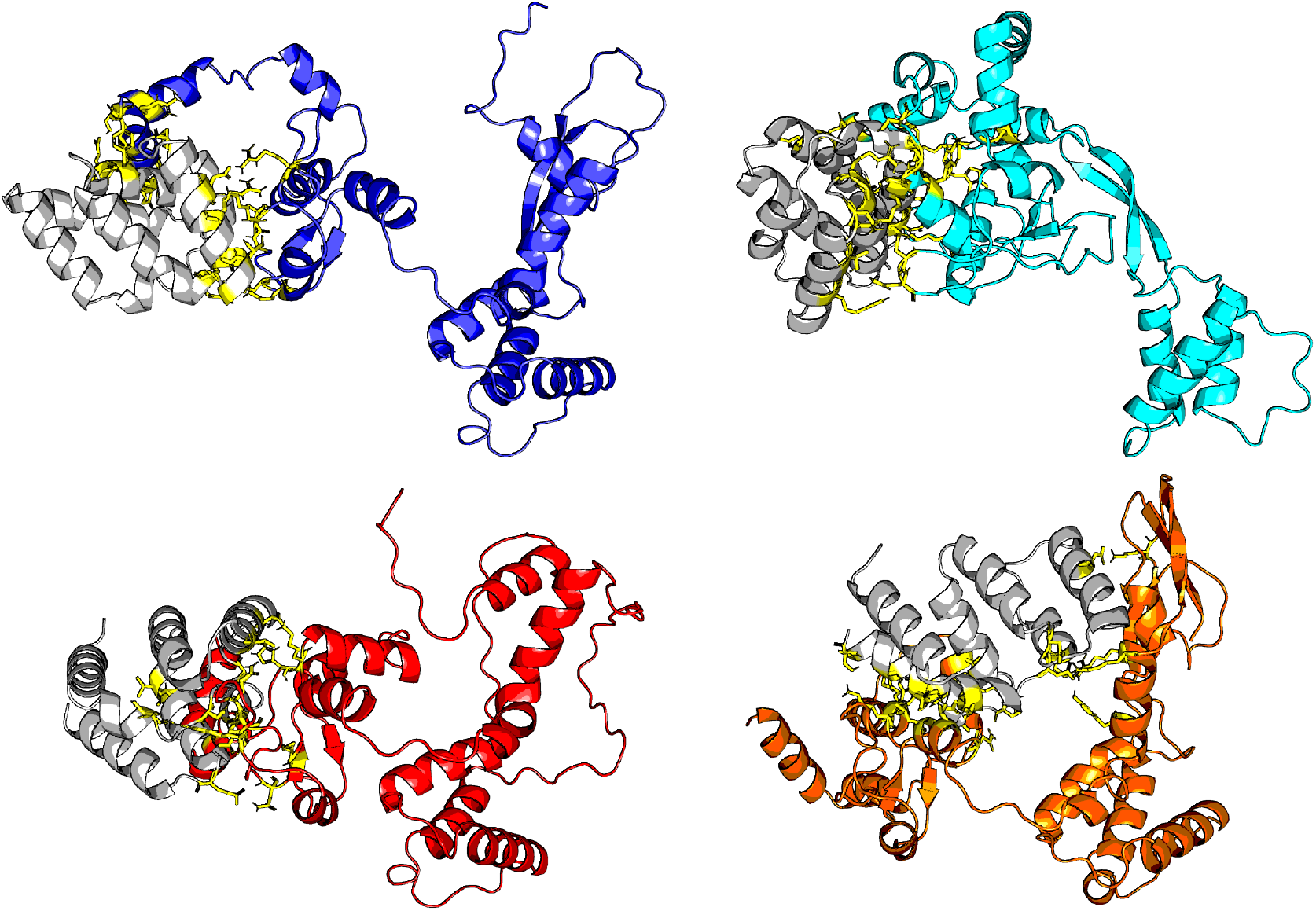
Docking results of MAVS (in grey) against PB2 structures. Each colored PB2 structure represents an individual virus [Blue: OQ958041 (A/American Wigeon/South Carolina/22-000345-001/2021, vaccine candidate), Cyan: PP577947 (A/Texas/37/2024), Red: PQ591825 (A/California/150/2024), and Orange: PQ809562 (A/Louisiana/12/2024)].

**Fig. 9.**
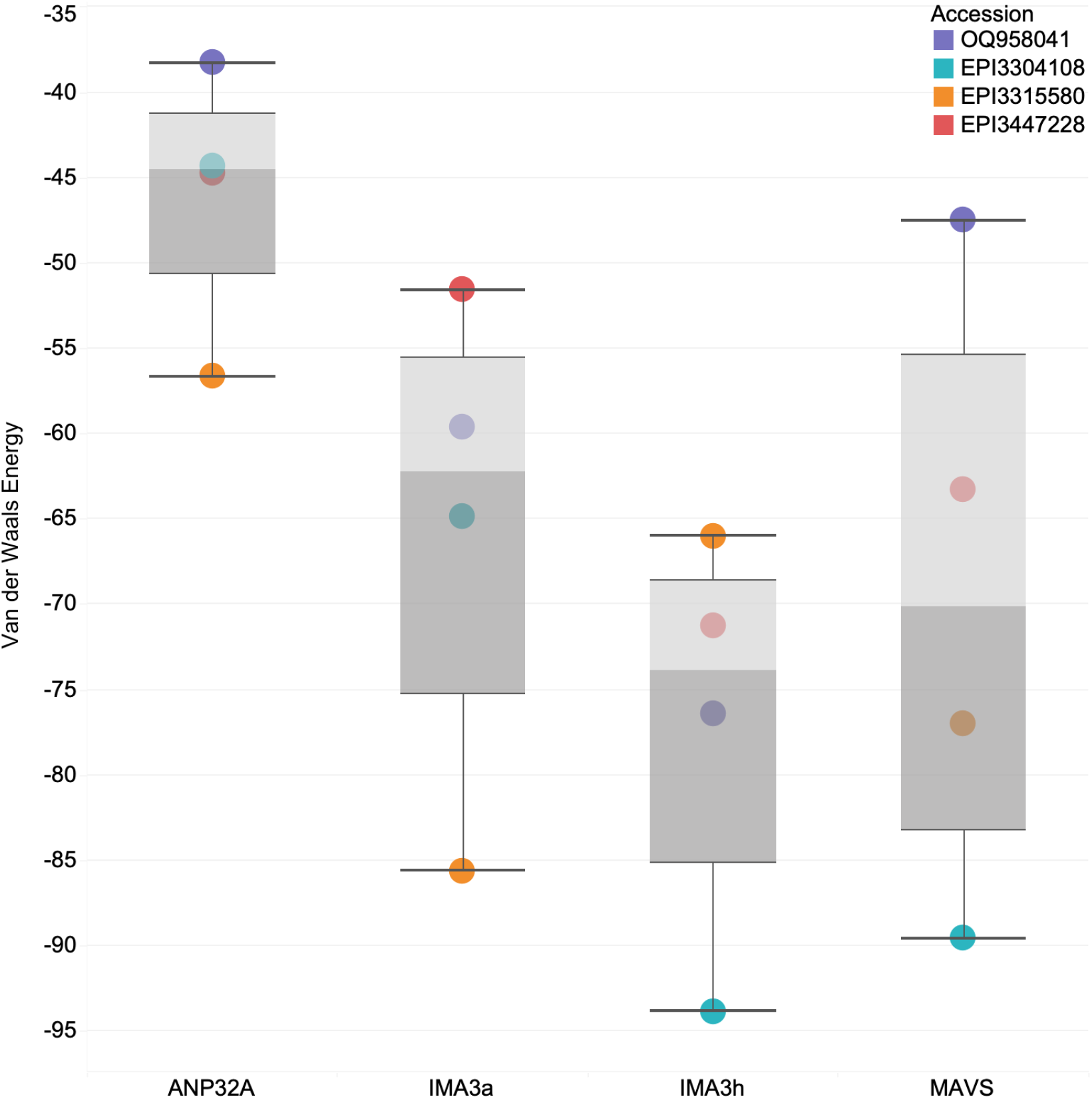
Boxplot graph showing the binding affinity, in Van der Waals’ energy on the y-axis, from docking simulations of Polymerase Basic 2 (with residue 588 mutations) against target proteins ANP32A, Importin-*α*3 (KPN4; avian and human variants), and Mitochondrial Antiviral Signaling (MAVS) on the x-axis. Lower Van der Waals’ energy values indicate stronger binding affinity. The viral sample Blue: OQ958041 (A/American Wigeon/South Carolina/22-000345-001/2021) represents the current vaccine candidate strain carrying the A588 residue. Other samples include Cyan: EPI3304108 (A/Great Blue Heron/AB/FAV-0505-25/2022) with the A588T mutation, Orange: EPI3315580 (A/American Crow/QC/FAV-0050-4/2022) with the A588S mutation, and Red: EPI3447228 (A/chicken/Nebraska/23-040358-001-original/2023) with the A588V mutation. Across all samples, binding affinity to ANP32A and MAVS remains like that of the vaccine candidate. Most of the 588 mutants show enhanced binding to human KPN4 compared to avian KPN4, consistent with adaptation to mammalian hosts; however, the A588S mutation (EPI3315580) uniquely shows a decreased affinity to human KPN4, suggesting a potential reduction in nuclear import efficiency. Although all mutant strains exhibit slightly weaker binding to human KPN4 than the vaccine candidate, the differences are minimal.

In addition to A588T/S/V, we checked the PB2-F6 constellation of mutations that are pointed out by Li et al., (2022) (54) to work in synergy with A588T to increase the pathogenicity and transmissibility in chickens and the virulence of mice of H10N8 in the single mutation of PB2-A588V (i.e. PB2: I292V, R389K, T598M, L648V, T676M). We report only wild-type genotypes for this constellation in our H5N1 isolates bearing A588T/S/V at this juncture. Moreover, these A588T/S/V bearing isolates are all currently at wild-type E627. However, many of the H5N1 viruses circulating in North America in 2022-24 carry 627K, including viruses hosted in animals (wild and domestic birds and mammals) and humans. Moreover, site 627 is under positive selection in our data.

Residue 676 exhibits mutations T676I, T676L, T676V, T676A. Mutation T676I underlies the expansion of lineages of H9N2, H7N7, H7N9 viruses in birds in the 2000’s and mammals and subsequently mutated to I676M (55). Caserta et al., (2024) (30) report the mutation T676A in genotype B3.13 H5N1 viruses along with other PB2 mutations (T58A, E362G, D441N, M631L). Despite these reports, the function of variants at site 676 has not been previously studied.

Structural Analysis of Commonly Used Antivirals Finally, we evaluated the binding of approved antiviral drugs to their respective protein targets from recent H5N1 isolates and from an older strain, A/chicken/Scotland/1959, as a comparison. In Figure 10, we see continued strength of binding affinity for most antivirals against their respective protein targets even in recent strains. Only in A/Louisiana/12/2024 (PQ809560) do we see a drastic decline in binding strength for antivirals Zanamivir (NA) and Baloxavir (PA).

**Fig. 10.**
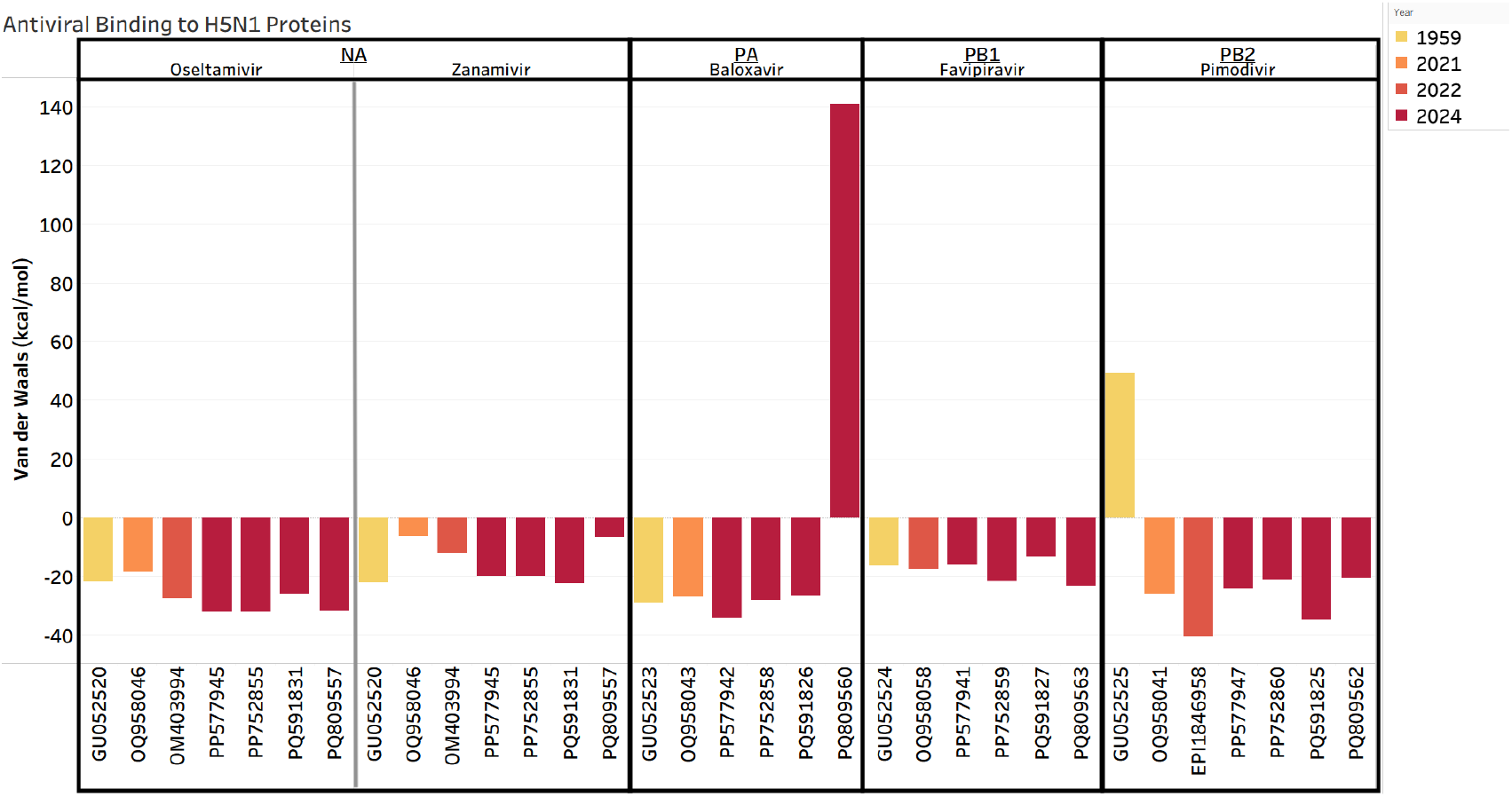
A bar plot showing the binding affinity metric (in Van der Waals’ energy; y axis) for each protein from each viral isolate to a respective antiviral (x axis). The viral isolates are labeled by their NCBI accession ID and refer to the strains A/chicken/Scotland/1959 (GU052520, GU052523, GU052524, GU052525), A/American Wigeon/South Carolina/22-000345-001/2021 (OQ958046, OQ958043, OQ958058, OQ958041), A/Astrakhan/3212/2020 (OM403994), A/cattle/Texas/24-009110-004/2024 (PP577945, PP577942, PP577941, PP577947), A/Texas/37/2024 (PP752855, PP752858, PP752859, PP752860), A/California/150/2024 (PQ591831, PQ591826,PQ591827, PQ591825), and A/Louisiana/12/2024 (PQ809557, PQ809560, PQ809563, PQ809562). Each divided panel shows the protein being represented by the accession IDs (NA, PA, PB1, and PB2) and the antiviral that it is binding to (Oseltamivir, Zanamivir, Baloxavir, Favipiravir, and Pimodivir). The color gradient from yellow to red signifies that year of collection for each isolate (ranging from 1959 to 2024).

## Discussion

There are five main points of discussion for analyses in this paper: 1) seasonality analysis of H5N1 infection cases across North America; 2) the structure of the clades of interest in North America and the natural selection they exhibit; 3) Natural selection and the changes of protein structure in HA leading to host promiscuity; 4) The changes in protein structure of PB2 increasing binding of host immune proteins; 5) The lack of mutations under selection for antiviral resistance and the maintenance of binding strength for most antiviral compounds in H5N1.

In this study, we combined phylogenetic, natural selection, and structural biology analyses to track the evolution of H5N1 clade 2.3.4.4b in North America and assess viral molecular adaptation to mammalian hosts. Additionally, we performed decomposition of seasonality to assess whether there are spikes of H5N1 infections across avian and mammalian hosts. Our findings indicate that recent viral lineages have evolved enhanced affinity for mammalian host receptors and immune factors, adaptations with significant public health implications. The integrated workflow presented here provides a rapid, computationally driven approach for moving from sequence data to functional predictions of viral risk that can be re-run as new data become available.

### Seasonality Trend

Our time-series analysis reveals a profound shift in the epidemiology of H5N1, characterized by an exponential increase in cases since 2021 and distinct, geographically-dependent seasonal patterns. The dramatic surge in cases corresponds directly with the introduction and subsequent spread of the highly pathogenic H5N1 clade 2.3.4.4b in North America. This clade has demonstrated unprecedented fitness, leading to widespread circulation in wild bird populations and a significant increase in spillover events into domestic poultry and mammals, which drives the exponential trend observed in our data (28).

The seasonal patterns identified in our analysis are likely driven by the migratory behavior of wild aquatic birds, the primary reservoir for influenza A viruses. The biannual peaks in May and November observed in avian populations outside the US align well with the timing of spring and autumn migrations along major global flyways (56). During these periods, large numbers of birds congregate at stopover sites, creating ideal conditions for efficient virus transmission and dispersal across continents (56, 57).

The annual peak in both avian and mammalian cases within the US during December suggests a slightly different dynamic. This late-year surge may reflect the congregation of migratory birds at their southern wintering grounds within the United States (58). Colder temperatures during this time can also enhance the environmental stability of the virus, further promoting transmission (58). Critically, the fact that mammalian cases peak concurrently with avian cases shows that the risk of spillover is directly coupled to this seasonal peak in viral infections of wild birds. These findings suggest that the late autumn and early winter months represent a predictable, high-risk period for H5N1 transmission in the U.S.

#### HA

The two main clades from the spread of H5N1 in North America in 2022-24 of interest in HA, are 1) a clade surrounding the human death in Louisiana (genotype D1.1 viruses) and 2) clade with human-bovine-avian transmission events (genotype B3.13). Notably, these clades of interest are phylogenetically distant in the HA tree. Moreover, these two HA clades have slightly different inter-host dynamics, as demonstrated by the character evolution and StrainHub analyses. The clade surrounding the human death in Louisiana has largely avian-primate exchanges. However, recent reports have indicated artiodactyl cases ((50); represented by Genbank PQ687463 in this subclade).

The clade in which human-bovine-avian interactions are emblematic shows high frequency of unidirectional transmission from Artiodactyla to Primates, bidirectional transmission between Galliformes and Primates, and unidirectional transmission from Carnivora to Primates. The mere fact that two phylogenetically distinct clades are currently infecting North American mammals demonstrates the non-canonical nature of the zoonotic evolution of HA. Moreover, these two clades have different selective regimes in the HA. The same motifs are under selection but not the same point mutations. The HA has two motifs of interest, the receptor binding domain (RBD; residues 113-265) and the stalk (residues 335-500). While both present antigenic properties, we focused on the RBD and its role for host selection via sialic acid conformation (59). The structural evolution of HA is towards viral host promiscuity. Our results show no binding strength differences for this HA motif for avian or mammalian receptors. This result is consistent with the hundreds of wild and domestic avian and mammalian hosts in North America (supplemental_table_1.csv and supplemental_table_2.csv).

For the HA subclade of interest for avian to human exchanges, including the lineage that led to the human death in Louisiana (Figure 4) we see the following sites of functional interest. Site 230 in the HA protein exhibits mutation A230V in our data. In the literature, this site is described in older H5N1 datasets as M230V. Site 230 is involved in immune escape and stabilization of the HA in experimental ferret infection (60).

For the HA subclade of interest for avian-bovine-human exchanges (Figure 5) we see the following sites of functional interest. Site 172 in the HA protein shows mutation A172T in our data. In the literature, A172T has been favored in vaccine candidate strains as the mutation confers increased replication properties in mammalian cell culture [e.g., Vero cells; (61).

Taken together, the distinct clades, each with wide but slightly different host dynamics and measurements of natural selection in HA, indicate that H5N1 (clade 2.3.4.4b) viral lineages have several non-canonical evolutionary and host pathways to become zoonotic emergent diseases of concern. The concerns include animal and human health and economic and nutritional well-being due to large agricultural losses (21). In each of the 100s of hosts now available to the virus, new mutational constellations and reassortment events can evolve leading to more concerning viral lineages.

Residues that have shown mutations under selection were used to define areas of interest of docking simulations. This resulted in the selection of the HA receptor binding domain and several domains within PB2 such as the amino terminal and the 627 domain (53). While mutations of interest were not directly involved as active binding residues in the docking simulations, these mutations were present near the active binding residues (Table 1). These mutations cause conformation changes, changes in charges and side chain sizes, and or effect residue polarity (62).

HA binding to SA remains a central determinant of host specificity. Previous studies have established that avian-adapted influenza strains preferentially bind *α*2,3-linked SA, whereas human-adapted strains favor *α*2,6-linked SA (63). Comparison of mutations within the HA RBD to active binding residues reveals that while the mutations are not directly involved in receptor binding, several sites that are under selection (e.g. 140, 147, and 230) lie within 10 amino acids of key binding sites. The proximity can introduce steric effects that influence binding affinity, altering host receptor specificity, and increasing host promiscuity. Dadonaite et al., (2024) (64) point out sets of mutations of interest they discovered in H5N1 HA in combined and computational and pseudoparticle experiments for receptor binding, HA stability, and immune escape mutations. We compared our mutations of interest from Table 1 with their mutations of interest with adjustments for numbering conventions. We did not find exact overlap in sites but we did fine overlaps in domains. While we saw no dramatic differences with the RBD of our selected isolates for the structural analysis. However, we observed a drastic difference in binding affinity of A/Louisiana/12/2024 HA protein binding to mammalian SA as compared to avian SA. This increase in preference to mammalian SA could signify a greater infection rate of mammalian host cells.

This promiscuity afforded by the adaptation of binding to a variety of receptors indicates that these strains retain the potential for widening zoonotic transmission and subsequent increases in mutation and reassortment events. Empirical studies have shown increased viral replication of H5N1 virus isolated from cattle in 2024 in a variety of mammalian cells including human and canid (65). The structural consistency in HA-SA interactions across different strains indicates that receptor binding alone may not be the primary barrier to human adaptation in these viruses. Additional factors such as polymerase function and host immune evasion are likely to contribute to host adaptation (66).

#### PB2

For PB2, we identified from our large dataset a single subclade of interest for zoonotic events. In our analyses of natural selection in this PB2 subclade, we find the following sites of functional interest. Site 249 in the PB2 protein exhibits many mutations in our dataset, including E249K. E249G, E249D, and E249V. Site 249 in the PB2 protein exhibits the E249K mutation in our data. H5N1 isolated from humans in Egypt in 2024 carried E249K (67). E249K was also carried in Human (H1N1) pdm09 cases in India in 2015 (68). A historical study of H5N1 viruses demonstrates that the E249G mutation improves replication in human cells (69). Most recently (2024) in H5N1 in dairy cows in Kansas the E249G mutation originated in mammals about this time in concert with mutations in NS1 (R21Q) and PB2 (E627K) (70). These mutations form a constellation associated with viral adaptation to mammalian hosts (70).

Similarly, E249D is among a constellation of mutations that led to increased pathogenicity of Eurasian avian-like swine lineage H1N1 (71). In H9N2, the I292V mutation improves virus replication in mammals (72). In 2024, a human traveler from India to Australia was infected with a recombinant (H5N1 clade 2.3.2.1a and 2.3.4.4b) virus with a clade 2.3.4.4b PB2 containing mutation V344M. V344M has been detected in wild birds and poultry in Asia since 2020 (73). Site 463 exhibits mutations I463M and I463V in our data. Mutation I463M in PB2 occurs in low frequency in viruses extracted from pigs experimentally infected with clade 2.3.4.4b H5N1 viruses derived from mink (74).

Results for position 588 are mixed. 1) H5N1 2.3.4.4b viruses carrying variation at 588 have only adapted a fraction of the genotypes for the constellation enabling viral promiscuity in H10N8 discovered by Li et al., (2022) (54). 2) Position 588 is under positive selection that is slightly statistically insignificant. 3) As site 588 has diverse results for binding ANP32A based on which genotype it carries (A588S caused a decrease in affinity to human proteins studied whereas A588T/V increased binding affinity to human proteins studied). At this juncture, variation at position 588 and its background are important to watch. Variants at position 588 could evolve into constellations already described and or variants at position 588 could evolve into a novel genomic constellation. The structural data indicates that some genotypes (e.g., A588T/V) are facilitating host promiscuity in the current H5N1 2.3.4.4b genomic backbone (Figures 7, 8, and 9).

Site 627 exhibits E627K and E627V mutations in our data. E627K is a well-studied mutation underlying the transmission of avian-hosted lineages H5N1 to mammals (75). E627K also increases the replication and virulence of the H5N1 in mammals (76). E627V allows many viruses of avian origin (H9N2, H7N9, and H3N8, which may share PB2 genes with H5N1 via reassortment) to infect and replicate in both chickens and mice by binding ANP32A proteins in both species (77). Experimental infection of mice with H5N1 virus bearing E627K leads to increased replication and mortality in mice (78). One human isolate in our data that bears E627K [Genbank: PP577947 A/Texas/37/2024(H5N1)] has been shown in experiments to transmit via respiratory droplets among ferrets, leading to mortality in most of the ferrets (79).

Taken together, these mutations (well-studied or understudied) point to the adaptation of H5N1 to increased transmission, virulence, and replication in a wide range of avian and mammalian hosts. Thus, our results on PB2 also support the hypothesis for the evolution of host promiscuity and increased viral replication to both avian and mammalian hosts in H5N1 clade 2.3.4.4b in North America in 2022-present. Misra et al., (2024) (80) note an array of 12 sites in PB2 that are of interest. We compared their list to ours and found that only sites 627 (which is well known) and 676 are in common.

PB2, a key component of the viral RNA polymerase complex, has been shown to influence host adaptation by interacting with cellular factors involved in nuclear trafficking and immune signaling (81, 82). Our docking analysis highlights strong PB2 interactions with MAVS and KPN4, particularly in recent strains, which bind much stronger to these proteins than the vaccine candidate (OQ958041) does. MAVS is a critical component of the hosts’ antiviral response, and an increased PB2 binding affinity enables immune evasion by hindering interferon signaling (43, 83). Moreover, Octaviani et al., (2025) (65) show a lack of RIG-1 levels in infected cells by 2024 viral strains. This result indicates antiviral signal antagonism by recent strains of the virus. RIG-1 is part of the innate immune response that causes cells to turn into an antiviral state and MAVS initiates this signaling pathway for RIG-1 (84). The results from Octaviani et al., (2025) (65) strengthen our findings as increased binding of MAVS impedes its signaling. Similarly, KPN4 and ANP32A mediate nuclear import of viral ribonucleoproteins. Improved binding between PB2 and KPN4 shows improved capability of nucleus localization in host cells to initiate influenza viral replication (44). Enhanced PB2-ANP32A interactions improve viral replication efficiency in mammalian cells as it has been shown to promote viral replication in Yu et al. (2022) (85).

Our results show a slight weakening of binding affinity of PB2 with ANP32A, the viral replication enhancer, in recent strains. However we see an increased binding affinity of PB2 with the nuclear transporter KNP4 and the immune response protein MAVS (i.e. in the A/Louisiana/12/2024 strain that caused a fatality as compared to the vaccine candidate strain).

#### NA, MP, PA and PB1

In our analysis of the NA, MP, PA, and PB1 proteins, we identified a limited number of amino acid sites under positive selection as shown in Table 2. For neuraminidase (NA), positive selection was detected at sites 16 and 81. While these sites are evolving, they are not located within or directly adjacent to the highly conserved active site of the neuraminidase enzyme, where inhibitors like oseltamivir and zanamivir bind. This suggests that the evolutionary changes at these positions are not likely to confer direct resistance to these drugs.

For the MP protein, positive selection was found at sites 61 and 88. However, resistance to adamantane-class MP inhibitors is primarily associated with mutations within the transmembrane domain and ion channel pore, most notably at site 31 (S31N) (86). The sites under selection fall outside this critical region, indicating that these mutations are unlikely to impact MP inhibitor effectiveness.

For the polymerase complex, positive selection was observed at sites 399 and 489 in the PA subunit. The primary resistance mechanism to the cap-dependent endonuclease inhibitor, baloxavir marboxil, involves a key mutation at site 38 (I38T) (12, 87). As the sites we identified are distinct from the known resistance-conferring mutations, their selection does not directly imply a loss of baloxavir efficacy.

The absence of any positively selected sites in the PB1 gene segment, as determined from our dataset of recent North American H5N1 isolates, is a significant finding. This result suggests that the PB1 protein, a core component of the RNA-dependent RNA polymerase, is currently under strong evolutionary constraint within this viral population to maintain its essential function. This high degree of conservation is particularly notable when contrasted with previous studies that have identified specific PB1 mutations associated with increased virulence and mammalian adaptation in other H5N1 lineages. For instance, substitutions like N105S have been shown to enhance polymerase activity in mammalian cells, contributing to increased pathogenicity (88). Furthermore, while resistance to the PB1-targeting antiviral favipiravir is rare in natural isolates, specific mutations such as K229R have been generated in vitro and can confer reduced susceptibility (89). The fact that our analysis did not detect positive selection at these or any other sites suggests that, within the evolutionary pressures currently acting on H5N1 in North America, adaptations in PB1 are not a primary driver of host-shifts or emerging antiviral resistance.

#### Antivirals

A crucial aspect of pandemic preparedness is understanding the continued effectiveness of existing medical counter-measures. Our positive selection analysis did not find any amino acid sites in respective proteins that lead to changes within their binding pockets for these antivirals. On the other hand, our structural analysis of antiviral binding provided insights into this question. We found that the binding affinity for most antivirals, including the widely used neuraminidase inhibitor oseltamivir, remained largely stable across recent H5N1 isolates compared to an older reference strain. However, we observed a notable decrease in the predicted binding affinity for Zanamivir (NA inhibitor) and Baloxavir (PA inhibitor) for the A/Louisiana/12/2024 isolate, which was associated with a fatal human infection. No public information is available regarding the type of treatment used on the patient, but CDC states primary treatment of Influenza infections with Oseltamivir (NA inhibitor) and then addition of Baloxavir for persistent infection (90). This finding highlights the potential for resistance to emerge on a strain-specific basis and could lead to decrease in treatment options. This discovery highlights the importance of continuous surveillance to monitor for increased spread of these resistant mutants (91). While our data support the continued primary use of oseltamivir in North America, the reduced predicted efficacy of other antivirals (i.e. Zanamivir and Baloxavir) against a highly pathogenic human isolate is a significant concern that warrants further investigation.

## Conclusion

Our results indicate that HA has achieved broad receptor-binding capability on both avian and mammalian cells, Moreover, we find that PB2 is undergoing adaptive changes that enhance innate immune evasion and replication in avian and mammalian hosts. This showcases the importance of monitoring both HA and PB2 in emerging influenza strains to assess their potential for human adaptation.

Our antiviral selection findings provide a mixed but largely reassuring picture regarding the potential for antiviral resistance in circulating H5N1 strains. While the virus continues to evolve in other regions of the proteins that are the main targets for current antivirals, our analysis did not identify positive selection at the primary amino acid sites known to be associated with resistance to the most widely used influenza antivirals.

Future research integrating structural modeling with functional assays will be crucial for validating these computational predictions and improving our understanding of influenza A H5N1 host adaptation mechanisms and potential therapeutics (14).

## Limitations

This study has limitations inherent to its computational nature. Our findings on binding affinity and selection are predictive and require empirical validation through in-vitro and in-vivo experiments. Furthermore, while we analyzed two key genes, HA and PB2, viral adaptation is a complex process involving the entire genome. Nevertheless, the workflow we have established—moving from large-scale phylogenetics to targeted structural biology—is a powerful tool for modern molecular disease surveillance. As sequence data becomes available, this approach can be rapidly deployed to assess new viral variants, providing early warnings and guiding public health responses before a crisis escalates.

## Data Availability

Raw sequence data, alignments and results from phylogenetics and analyses of selection, predicted protein structures, and protein-protein binding data are available on GitHub at https://github.com/colbyford/Influenza_A_2.3.4.4b_Analyses.

## Code Availability

Code for running the analyses used in this study is available on GitHub at: https://github.com/colbyford/Influenza_A_2.3.4.4b_Analyses

## Author Contributions

DJ built the datasets, conducted protein nucleotide sequence analysis, analysis of character evolution, conceived of the hypotheses and means of testing. KO conducted analysis of selection. SGM and CTF conducted analyses of structural biology and protein-protein interactions. KO and RA managed data. All authors wrote and edited the manuscript.

## Acknowledgments

We acknowledge all GISAID data contributors (i.e., the authors and their originating laboratories) responsible for obtaining the specimens, and their submitting laboratories for generating the genetic sequence and metadata and sharing via the GISAID Initiative, on which this research is based. We acknowledge the following entities at the University of North Carolina at Charlotte: Academic Affairs, The Office of Research (Ignite Award), The Center for Computational Intelligence to Predict Health and Environmental Risks (CIPHER), The Department of Bioinformatics and Genomics, The College of Computing and Informatics, and the University Research Computing group. We gratefully acknowledge the support of the Belk Family and the Levine Scholars program.

